# Circadian regulation of *MGMT* expression and promoter methylation underlies daily rhythms in TMZ sensitivity in glioblastoma

**DOI:** 10.1101/2023.09.13.557630

**Authors:** Maria F. Gonzalez-Aponte, Anna R. Damato, Laura Lucía Trebucq, Tatiana Simon, Sandra P. Cárdenas-García, Kevin Cho, Gary J. Patti, Diego A. Golombek, Juan José Chiesa, Erik D. Herzog

## Abstract

**Background:** Glioblastoma (GBM) is the most common primary brain tumor in adults. Despite extensive research and clinical trials, median survival post-treatment remains at 15 months. Thus, all opportunities to optimize current treatments and improve patient outcomes should be considered. A recent retrospective clinical study found that taking TMZ in the morning compared to the evening was associated with a 6-month increase in median survival in patients with *MGMT*-methylated GBM. Here, we hypothesized that TMZ efficacy depends on time-of-day and O^6^-Methylguanine-DNA Methyltransferase (*MGMT*) activity in murine and human models of GBM.

**Methods and Results:** *In vitro* recordings using real-time bioluminescence reporters revealed that GBM cells have intrinsic circadian rhythms in the expression of the core circadian clock genes *Bmal1* and *Per2*, as well as in the DNA repair enzyme, *MGMT*. Independent measures of *MGMT* transcript levels and promoter methylation also showed daily rhythms intrinsic to GBM cells. These cells were more susceptible to TMZ when delivered at the daily peak of *Bmal1* transcription. We found that *in vivo* morning administration of TMZ also decreased tumor size and increased body weight compared to evening drug delivery in mice bearing GBM xenografts. Finally, inhibition of MGMT activity with O^6^-Benzylguanine abrogated the daily rhythm in sensitivity to TMZ *in vitro* by increasing sensitivity at both the peak and trough of *Bmal1* expression.

**Conclusion:** We conclude that chemotherapy with TMZ can be dramatically enhanced by delivering at the daily maximum of tumor *Bmal1* expression and minimum of *MGMT* activity.

**Graphical Abstract:** 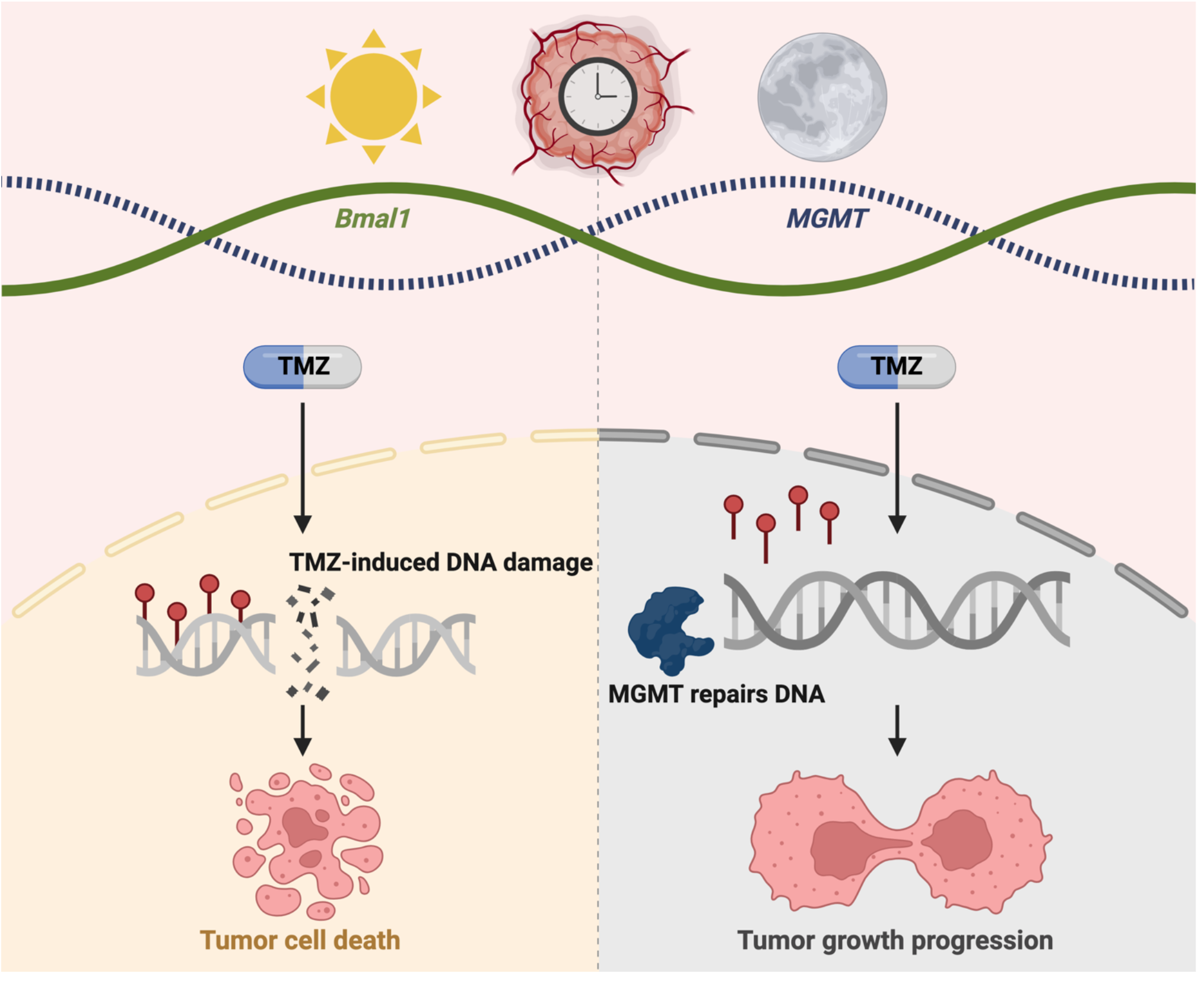

## Introduction

Gliomas are the most common brain malignancies, consisting largely of cells that resemble astrocytes, oligodendrocytes, oligodendrocyte precursor cells, and earlier neural stem cells [1], [2]. Glioblastoma (GBM) is the most common and aggressive glioma in adults, accounting for 54% of all gliomas, and 16% of all primary brain tumors [3]. In the United States, 12,000 adults are diagnosed annually at a median age of 64 years [4]. The current standard of care for GBM consists of maximal safe surgical resection, followed by radiation and chemotherapy with the DNA alkylator, Temozolomide (Temodar^®^, TMZ), and tumor-treating fields [5], [6]. When introduced into the standard of care for GBM approximately 20 years ago, TMZ extended median survival by 2.5 months, which was heralded as a dramatic improvement in treatment [7], [8]. In clinical applications, however, progression-free survival at 6 months in patients receiving TMZ was only 46% and even lower for recurrent GBM (17%) [9]. Despite extensive research and efforts to improve outcomes, median survival time remains approximately 15 months, and 5-year survival is less than 5%, after diagnosis [3]. Thus, the importance of further research to optimize current, and develop new, treatments against GBM remains highly significant and all avenues to lengthen survival should be pursued.

Among first line chemotherapy drugs used to treat GBM patients, TMZ has many advantages including oral administration, easy penetration through the blood-brain barrier, and no known toxic interactions with other drugs used in the clinic [10]. TMZ is a DNA alkylating agent that triggers the death of GBM cells by attaching a methyl group to purine bases of DNA (O6-guanine; N7-guanine and N3-adenine) [10]. The most common TMZ-induced cytotoxic lesion is O6-methylguanine (O6-MeG), which can be removed by the DNA repair enzyme O6-Methylguanine-DNA Methyltransferase (MGMT) in tumors that express this protein [10]. Expression of this DNA repair enzyme confers resistance to chemotherapy and represents a challenge in treating patients with unmethylated *MGMT* promoter sequences [10], [11]. In contrast, patients with *MGMT*-promoter methylated tumors (i.e., silenced *MGMT*) respond better to chemotherapy and have a better prognosis. In a retrospective study of morning versus evening TMZ, efficacy was greater in patients with *MGMT*-methylated tumors, who exhibited a longer median survival of 6 months when receiving morning TMZ compared to evening [12]. Thus, *MGMT* expression and activity confers resistance to TMZ in patients and may vary with time of day in GBM.

Circadian rhythms in gene expression and physiology are ubiquitous across cell types and species [13], [14]. Unlike some other cancers where circadian rhythms tend to be disrupted, well-studied GBM models have reliable circadian rhythms in gene expression [15]–[17]. Furthermore, expression of the MGMT protein has been shown to oscillate daily in healthy mouse liver cells, peaking during the subjective night (Circadian Time, CT19) [18]. It is unknown whether *MGMT* expression varies with time of day in GBM cells. Previous research has found that primary isolates from GBM patients and immortalized human and murine GBM cell lines, have high-amplitude daily rhythms in *Per2* and *Bmal1* clock gene expression, and in sensitivity to chemotherapy *in vitro* [15]. Thus, GBM biology and response to chemotherapeutic agents can vary with time of day. However, we do not know the mechanisms driving daily rhythms in TMZ efficacy or whether these daily rhythms in sensitivity are conserved across the diversity of gliomas. Several additional factors that could influence TMZ efficacy in the clinic also vary with time of day. For example, absorption and excretion of drugs in the blood varies by time of day, as does permeability of the blood-brain barrier [19]–[21]. Activation of checkpoint kinases that trigger apoptosis after induced DNA damage has also been shown to change based on time of day via an interaction with the clock genes *Per1* and *Per3* [22]–[24]. It is unknown if drug resistance in GBM can be ameliorated through strategically timed drug delivery to maximize tumor death while minimizing side effects.

Here, we aimed to maximize tumor cell death by targeting circadian molecules in GBM models. We found that human and murine GBM cell lines exhibit daily rhythms in TMZ sensitivity with the highest efficacy observed at the peak of *Bmal1* and trough of *MGMT in vitro*, and morning administration *in vivo*. This correlates with daily rhythms in *MGMT* methylation that peak before its transcription. Inhibiting MGMT activity *in vitro* enhances TMZ sensitivity as a function of circadian time. We conclude that TMZ efficacy and GBM outcomes can be improved by targeting daily rhythms in *MGMT* expression and activity.

## Results

### GBM sensitivity to TMZ increases around the daily peak of *Bmal1* expression

To test whether models of GBM differ in their sensitivity to TMZ over the day, we transduced human LN229 and murine GL261 cells with luciferase reporters of *Bmal1* or *Per2* transcriptional activity (B1L or P2L, respectively). In human LN229 cells, we found intrinsic daily rhythms in *Bmal1* and *Per2* expression over 72 h in culture, with *Bmal1* and *Per2* peaking in antiphase at CT4 or CT16, respectively (Fig. 1a, all recordings had cosine fits with correlation coefficients, CC>0.9). We next treated cultures with TMZ at one of eight circadian phases of *Bmal1* expression and found 100µM TMZ was more toxic to LN229 cells at the daily peak of *Bmal1* (Fig. 1b, *p=0.03; 72 h after a single-dose TMZ treatment, average trace scored circadian by JTK cycle p < 0.05). Specifically, we found that 21% of LN229 cells survived TMZ treatment around peak *Bmal1*, while 69% survived when treated at the trough. This time-of-day sensitivity reproduced in three different LN229 and GL261 biological replicates (Fig. 3). TMZ dose-dependently killed LN229 cells, dependent also on circadian time of administration (Supplementary Fig. 1, IC50 at CT4=195µM vs. IC50 at CT12= 321µM). The daily rhythm in TMZ sensitivity was not detectable at 10, 200 or 1000µM TMZ (Fig. 1b, p>0.05, average traces not scored circadian by JTK cycle p > 0.05). Together, these results demonstrate that an intermediate dose of TMZ aligned to the peak of *Bmal1* in GBM cells results in greater cell death.

**Fig. 1.**
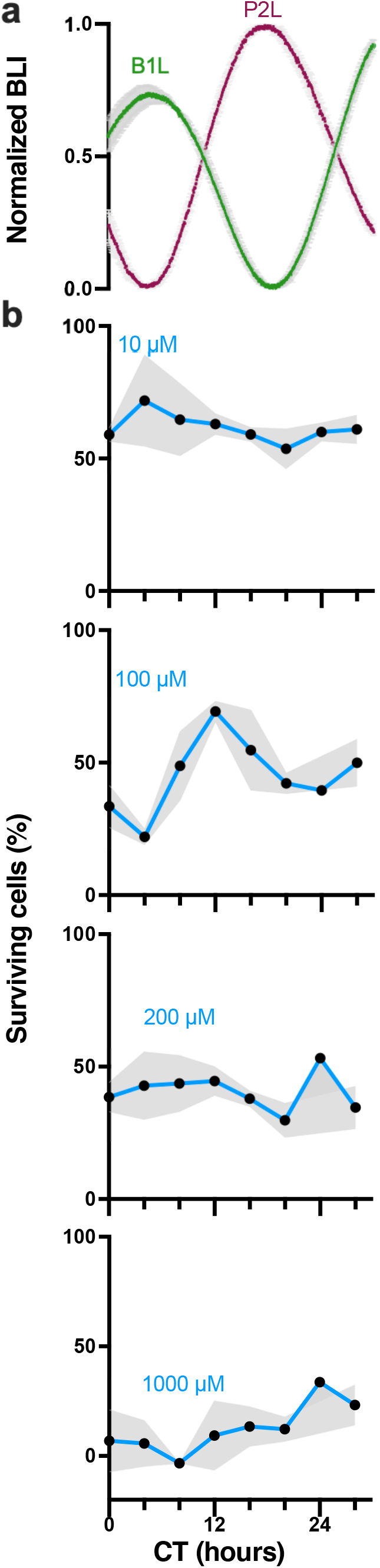
TMZ maximally inhibits growth at the daily peak of *Bmal1* expression in GBM cells *in vitro*. **a)** Circadian rhythms in clock gene expression recorded as bioluminescence from human LN229 GBM cells transduced with a *Per2-* or *Bmal1*-driven luciferase reporter (P2L or B1L, n=2 cultures each, mean ± SEM). **b)** Survival of LN229-B1L cells varied with the circadian time (CT) and dose of TMZ treatment *in vitro*. For example, more cells survived 100µM TMZ when administered around the trough of *Bmal1* (CT12) than around the peak (CT4). Higher doses of TMZ yielded more cell death and less circadian dependence (n=3 cultures per dose at each time, mean ± SEM, 100µM JTK cycle p = 0.001, ns for 10, 200, 1000µM)

To evaluate *in vivo* TMZ sensitivity as a function of time of day, we implanted immunocompromised nude mice with human LN229 cells, and C57 mice with murine GL261 cells (Fig. 2a). These orthotopic GBM models expressed either circadian (*Per2*-luciferase) or constitutive (*Ef1α*-luciferase) reporters to monitor tumor growth as bioluminescence after implantation. We delivered 100mg/kg TMZ or vehicle (10% HPMC) by oral gavage in the morning (11:00 a.m., 4 h after daily light onset) or evening (6:00 p.m., 1 h before daily light offset) for five days starting 11-13 days after implantation. We found no significant difference in tumor growth between mice treated with vehicle at each time point tested, and thus data were combined. Further, we found no significant difference in bioluminescence counts between tumors expressing *Per2*:luc or *Ef1α*:luc (Fig. 2b-d), suggesting that both reporters can be used to track tumor growth *in vivo*. We found that tumor bioluminescence declined faster in mice treated with TMZ in the morning, compared to evening and vehicle treatment regardless of the reporter (Figs. 2b and c, ****p < 0.0001 in LN229*-Ef1α*:luc, *p = 0.05 in LN229-P2L) or cell line (Fig. 2d, *p = 0.03). In parallel, we found that body weight increased for mice treated with TMZ in the morning but decreased in mice treated with TMZ in the evening or with vehicle (Fig. 2e-g, *p = 0.02 in LN229*-Ef1α*:luc, *p = 0.01 in LN229-P2L, *p = 0.02 in GL261*-Ef1α*:luc). Together, these data demonstrate that TMZ reduced GBM tumor size and slowed disease progression more when administered in the morning, compared to evening, in both human and murine models of GBM.

**Fig. 2.**
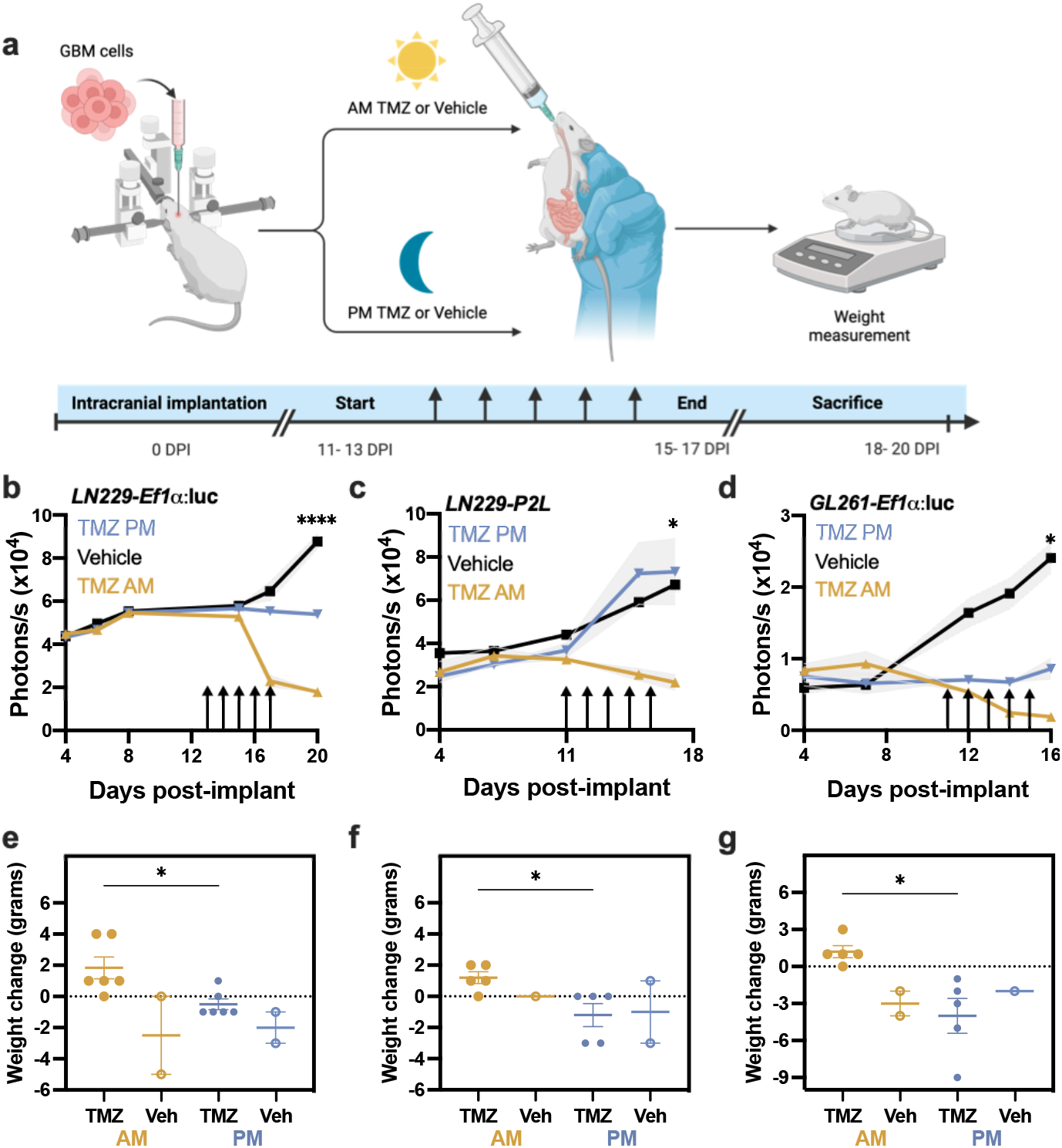
GBM xenografts show elevated sensitivity to TMZ in the morning, compared to evening. **a)** Experimental paradigm testing TMZ chronotherapy in GBM orthotopic xenograft models. **b)** LN229-Ef1α implants grew less following five daily doses (arrows) of 100mg/kg TMZ in the morning (ZT4, mean ± SEM) compared to those treated with TMZ in the evening (ZT11) or vehicle (****p < 0.0001 AM vs. PM at 20 days post-implantation, DPI). We quantified tumor size as total bioluminescence at each 11am measurement. **c)** Mice bearing LN229-P2L xenografts also showed less tumor growth when treated in the morning compared to vehicle or evening TMZ (TMZ PM, *p < 0.05 AM vs. PM at 17 DPI). **d)** Similarly, GL261-Ef1α xenografts grew less when treated in the morning compared to vehicle or evening TMZ (*p < 0.05 AM vs. PM at 16DPI). **e-g)** Mice bearing LN229 or GL261implants treated with TMZ in the morning lost less weight from start to the end of TMZ treatment compared to those treated in the evening or with vehicle (mean ± SEM; *p < 0.05)

We also tested TMZ at other times of day, doses, and disease state (Supplementary Fig. 2a-b). Mice were implanted with GL261-P2L or LN229-P2L cells and, immediately after implantation (4 days post-implant), started receiving 70mg/kg TMZ by oral gavage for two weeks, 5 days on and 2 days off, at either 8:00 a.m. or 7:00 p.m. (Zeitgeber Time, ZT1 or ZT12). After a total of 10 treatments with a lower TMZ dose, we found no significant difference between groups (Supplementary Fig. 2a-b, p>0.99). These results indicate that time-dependent TMZ sensitivity depends on disease progression status at start of treatment, dose, and circadian time.

### Time-dependent sensitivity to TMZ does not depend on blood-brain barrier penetration

Unlike molecules that are actively transported across the blood brain barrier, and thus subject to daily rhythms in efflux transporter activity, TMZ passively diffuses [25]. We quantified TMZ and its catabolite, 4-amino-5-imidazole-carboxamide (AIC), via mass spectrometry after morning or evening gavage (70mg/kg) in mice. We found similar levels of TMZ and AIC in blood and brain after morning or evening TMZ delivery (Supplementary Fig. 3a-d). Consistent with prior reports and modeling [25], [26], TMZ and AIC levels were lower in brain than blood and rapidly declined within 4 h (****p < 0.0001). We conclude that time-of-day differences in response to TMZ are likely due to tumor-intrinsic circadian rhythms and not differences in drug distribution.

### Daily rhythms in *MGMT* expression determine phase-dependent TMZ sensitivity

To test the hypothesis that TMZ sensitivity relates to circadian regulation of the DNA repair enzyme MGMT, we first assessed whether GBM cells exhibit daily rhythms in *MGMT* gene expression. Analysis of the *MGMT* promoter sequence (NG_052673, -2591 bp to +123 bp) yielded a total of 12 E-box sequences (CACGTG), highlighting the potential for transcription of this gene to be modulated by the clock protein *BMAL1*. To record *MGMT* gene activity in real-time, we transduced GBM cell lines with an *MGMT*-luciferase reporter (*MGMT*:luc, courtesy of Dr. Markus Christmann [27]) and recorded its transcription every 3 min for 48 h. We found daily rhythms in *MGMT*:luc expression in LN229 GBM cells *in vitro*, with expression peaking around CT12, corresponding to the trough of *Bmal1* (Fig. 3a, CC>0.9). We next performed qPCR and qMSP on LN229 cells collected every 4 h over 48 h. We found daily rhythms in *MGMT* gene expression and methylation status, with *MGMT* transcription peaking at CT12, and methylation being at its lowest point at CT8 (Fig. 3b-c, average trace scored circadian by JTK cycle, p < 0.05), consistent with the *MGMT*:luc reporter and indicating circadian regulation of DNA repair.

**Fig. 3.**
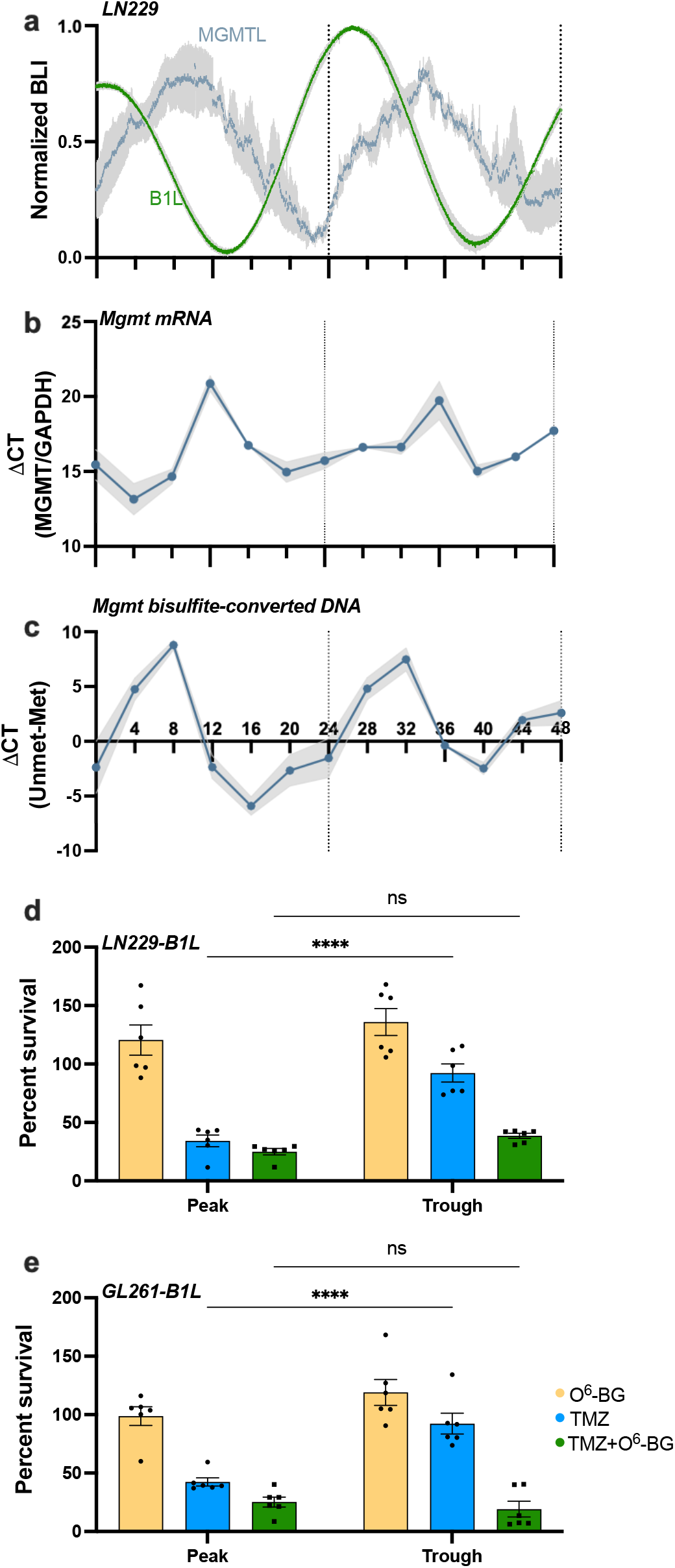
TMZ sensitivity depends on MGMT activity and correlates with daily rhythms in *MGMT* mRNA abundance and promoter methylation *in vitro*. **a)** Circadian rhythms in *MGMT* expression peak approximately 10 h after peak *Bmal1* expression in human LN229 GBM cells. Bioluminescence of cells transduced with either *MGMT*-luciferase (MGMTL) or *Bmal1*-luciferase (B1L) was recorded for 48 h *in vitro* (n=5 cultures each, mean ± SEM). **b)** LN229 cells collected every 4 h over 48 h show daily rhythms in *MGMT* mRNA (n=6 cultures each, mean ± SEM, JTK cycle p = 0.05) and **c)** *MGMT* promoter methylation (n=3 cultures each, mean ± SEM, JTK cycle p = 0.002). **d)** LN229-B1L and **e)** GL261-B1L cells show decreased cell number when treated with 100µM TMZ at either the peak or trough of *Bmal1*, with highest sensitivity at the peak. This time-of-day-dependent sensitivity is abrogated when co-treating with the MGMT inhibitor O^6^-BG (n=6 per group; mean±SEM; ****p < 0.0001, ns indicates p > 0.05)

To test if daily rhythms in TMZ efficacy depend on MGMT activity, we treated LN229-B1L and GL261-B1L cultures with 100 µM TMZ and the MGMT inhibitor, O^6^-benzylguanine (20µM O^6^-BG) or vehicle. As in previous experiments, we found that the day-night difference in TMZ was abrogated such that TMZ killed more cells at the peak of *Bmal1* (42% survival for GL261; 34% survival for LN229 cells) than at the trough (92% survival for both GL261 and LN229 cells) (****p < 0.0001), but not when co-delivered with O^6^-BG (Fig. 3d-e, p>0.05). Together, these results demonstrate that daily rhythms in TMZ sensitivity in GBM cells depend upon circadian *MGMT* expression. We conclude that chemotherapy with TMZ can be dramatically enhanced by delivering when *Bmal1* expression in the tumor reaches its daily maximum and *MGMT* transcription is at its minimum.

## Discussion

### TMZ efficacy against GBM can be enhanced by delivery at the best time of day

The results presented here demonstrate that delivering chemotherapy at a specific time of day can significantly decrease tumor growth and improve disease outcomes for human and mouse models of GBM. Since the FDA approval of TMZ in 1999 and tumor-treating fields in 2011, no new treatments have been introduced into the standard of care for GBM and, even with aggressive therapy, patient prognosis and survival remain very poor. Here we reveal daily rhythms in sensitivity to chemotherapy modulated by the daily expression of the DNA repair enzyme *MGMT*. If validated in patients, these findings can be rapidly translated to prescribe TMZ and *MGMT* inhibitors at the best time of day for GBM patients. Taken together, these data support the potential for TMZ to be chronomodulated and identify a targetable mechanism promoting resistance to timed chemotherapy.

Our results reveal daily rhythms in sensitivity to TMZ in well-studied human and murine GBM models. We found a 3-fold reduction in GBM tumor bioluminescence with morning versus evening TMZ *in vivo*. This correlates with the 4-fold greater tumor suppression when treating GBM cells at the peak (CT4) of daily *Bmal1*, and minimum of daily *MGMT* expression, *in vitro*. The changes in body weight further suggest that morning TMZ slowed disease progression, compared to evening treatment. Consistent with these findings, a recent retrospective clinical study found that patients receiving TMZ in the morning had an increased median survival of 6 months, compared to those receiving it in the evening [12]. Based on the convergent evidence for daily rhythms in sensitivity to TMZ in various cellular and orthotopic models of glioma and in human GBM patients [15], we conclude that TMZ efficacy can be enhanced by administering it in the morning and in alignment with low *MGMT* and high *Bmal1* gene expression.

One caveat to these results is that we do not know if all GBM tumors synchronize to the host in the same way and thus, if daily *MGMT* expression and TMZ sensitivity vary depending on tumor or host genetics. Because GBM is a highly heterogenous disease, it is possible that intrinsic clock gene expression, synchronization of circadian rhythms to the host, and time of peak sensitivity to TMZ vary depending on tumor type, status, localization, and the patient’s individual circadian rhythm. Thus, to incorporate TMZ chronotherapy into the standard of care for GBM patients, it will be important to understand when *MGMT* and *Bmal1* peak in individual human GBM tumors, whether this varies between patients, and the optimal dose and time required to obtain maximum drug effects.

### Circadian regulation appears to be common to a variety of GBM models

A broad range of GBM cells and GBM stem cells have been found to exhibit intrinsic circadian rhythms in clock gene and protein expression, and in sensitivity to therapeutic drugs [15]–[17]. Our data further expands on these results revealing 24-hour rhythmic expression profiles in *Per2* and *Bmal1* transcription in murine (GL261) and human (LN229) GBM models. We found that *Bmal1* and *Per2* transcription peak at similar subjective times of day in GBM *in vitro* compared to the SCN, prefrontal cortex, hippocampus, amygdala, among other tissues [28]. Consistent with previous studies [15], we found daily rhythms in sensitivity to TMZ with the highest efficacy observed at the peak of *Bmal1 in vitro* and in the morning *in vivo*. While studies have shown circadian regulation of TMZ *in vitro*, there has been no mechanistic exploration as to why efficacy of this chemotherapeutic drug varies with time of day. Here we found circadian rhythms in the transcription and promoter methylation of the DNA repair enzyme *MGMT* that peak in anti-phase to *Bmal1* expression. *MGMT* peak expression was found to be at CT12, consistent with previous studies that have described daily rhythms in *MGMT* expression in mouse liver tissue [18]. This daily anti-phase variation in *Bmal1* and *MGMT* expression suggests that the circadian clock may modulate rhythms in DNA repair and thus, provides a time-sensitive window when TMZ can induce greater DNA damage without competing with MGMT. This model is supported with results showing that inhibiting MGMT activity with O^6^-BG abrogates time-dependent TMZ sensitivity. Altogether, our findings suggest a potential mechanism modulating daily sensitivity to chemotherapy and introduce the concept of combination therapy in which MGMT inhibition is combined with TMZ to maximize efficacy and increase tumor death.

### Determining MGMT promoter methylation status may vary with time of day of GBM sample collection

One surprising finding is that methylation of the *MGMT* promoter varies with time of day in LN229 cells, a GBM model widely considered to be unmethylated. We found low *MGMT* methylation levels at CT8, and high at CT16, suggesting an alternative mechanism by which daily rhythms in *MGMT* expression are modulated by post-translational modifications that silence or activate its transcription. Importantly, *MGMT* promoter methylation is a strong prognostic factor in the therapy of GBM patients [29]–[32]. Methylation of the *MGMT* promoter is associated with a better response to TMZ chemotherapy and longer median survival [31]. However, our results suggest that determining a tumor’s methylation status might vary depending on the time of day at which the sample is collected. Moreover, one study found changes in *MGMT* promoter methylation status from primary tumor to relapse in some GBM patients [33]. This variation could be explained by daily rhythms in promoter methylation and poses the question of whether daily rhythms in *MGMT* methylation change with disease progression or after aggressive therapy.

We conclude that circadian rhythms in gene expression and response to therapies are highly reproducible and conserved across multiple GBM models. We present two potential mechanisms modulating timed sensitivity to TMZ and highlight the importance of circadian regulation in both the diagnosis and treatment of GBM. It remains unknown whether daily rhythms persist *in vivo*, if they synchronize to the hosts circadian rhythm, and whether circadian signals entrain rhythms in GBM to coordinate gene expression between the tumor and the body. Our results set the stage for future studies in tumor synchrony to the host, circadian medicine, and studies that translate to clinical care by using human cell lines and human data. These studies will need to account for details such as tumor location, sex of the patient, available sequencing data, and times of surgery or tissue fixation. Future work will elucidate whether circadian rhythms in GBM can be leveraged to improve current therapies and if personalizing treatments based on a patient’s intrinsic circadian rhythm can prolong their survival.

### Implementation of chronotherapy into the standard of care for GBM requires no additional approvals and is consistent with recent retrospective chart study

While largely understudied for TMZ and GBM, the concept of chronotherapy (i.e., the practice of considering time of day in treating a disease) has been shown to improve outcomes in several cancers, including acute lymphoblastic leukemia, colorectal, ovarian, and other gynecological cancers [20], [34]–[36]. Importantly, novel therapeutic drugs for GBM, such as the inhibitor of Rac family small GTPase 1, 1A-116, showed daily rhythms in efficacy *in vitro* and *in vivo* [17]. In addition to tumor sensitivity to TMZ, in a phase I clinical trial administering TMZ at morning or evening times, adverse events were equally common in both patient groups, indicating that host side effects to TMZ are not different at the two administration times tested [37]. Incorporating chronotherapy into the standard of care for GBM patients requires no additional approvals or clinical trials, as opposed to the development of new therapies that can take decades. Our data are consistent with a previous study that associated longer patient overall survival with morning TMZ administration [12]. Because TMZ is taken orally at home, translation of these findings to patients is relatively simple. Future studies should focus on evaluating circadian medicine personalized to the tumor type and daily sleep patterns of the patient in addition to controlling for factors such as age, sex, socioeconomic status, and time since diagnosis.

## Methods

### Cell culture, lentiviral transductions, and bioluminescence recordings i*n vitro*

LN229 and Glioma 261 (GL261) cells were grown in DMEM (Gibco) or RPMI-1640 (Sigma-Aldrich), respectively, supplemented with FBS (Fisher) and 1% Pen/Strep (Thermo Fisher). Cells were grown in a 37*°*C incubator with a 5% CO2 environment. Passage number in all experiments ranged from four to twelve. GBM cells were transduced with lentiviral reporters expressing firefly luciferase driven by the mouse *Bmal1* (*Bmal1*-luc [38], [39]), *Period2* (*Per2*-luc [40]), *Ef1α* (*Ef1α-*luc, obtained from GenTarget Inc.), or O^6^-Methylguanine-DNA methyltransferase (*MGMT*-luc [27]). After 24h of transduction, infected cells were selected using blasticidin (1.25 μg/mL, Thermo Fisher). Luciferase expression was confirmed by recording bioluminescence as photons per 180 seconds *in vitro*. Cells were cultured in media containing 0.1mM D-luciferin (Goldbio) and placed under photomultiplier tubes (PMTs) (Hamamatsu Photonics). See Extended Materials and Methods.

### *In vitro* cell growth assays and pharmacology

To assess daily TMZ sensitivity, cells were plated and synchronized via serum shock with 50% FBS for 2hrs, followed by a media change, every 4hrs so that at one treatment time, plates spanned 0-, 4-, 8-, 12-, 16-, 20-, 24-, and 28-hrs post-serum shock (HPS). Cells were treated with one of four TMZ concentrations or vehicle (DMSO, 0.2%). To assess whether daily rhythms in TMZ sensitivity depend upon MGMT activity, cells were treated with 20μM of the MGMT inhibitor O^6^-BG or vehicle (DMSO, 0.2%) at two different time points. In all experiments, cells were fixed after 72hrs with cold methanol and stained cells with 4′,6-diamidino-2-phenylindole (DAPI, 2μg/mL). DAPI fluorescence was quantified with the Infinite 200 PRO plate reader (V_3.37_07/12_Infinite, Tecan Lifesciences). See Extended Materials and Methods.

### Quantitative real-time PCR (qRT-PCR) and Methylation Specific PCR (qMSP)

RNA was extracted from LN229 cells collected every 4hrs for a period of 48hrs using TRIzol reagent. RNA was purified and converted to cDNA by RT-PCR. For qMSP, bisulfite DNA conversion was performed as previously described. Gene expression changes were further probed using iTaq™ Universal SYBR® Green Supermix (Bio-Rad). All procedures were done in triplicate in two biological replicates. The primer sequences for qRT-PCR and qMSP are listed in Table S1 and S2, respectively. See Extended Materials and Methods.

### Orthotopic xenografting

200,000 GBM cells were stereotactically implanted into the right caudate putamen of 10-week-old male and female NU/J (The Jackson Laboratory, Strain #002019) for human models, or C57BL/6J (The Jackson Laboratory, Strain #000664) for murine models. Following surgery, mice were housed in individual cages, monitored, and treated with analgesic for three days post-implant. Cells were allowed to engraft for 7 days before performing *in vivo* bioluminescence imaging to measure tumor size.

### *In vivo* bioluminescence imaging

Mice were housed in standard 12h light/12h dark conditions in wheel-cages to record locomotor activity in one-minute bins. Following orthotopic xenografting, tumor size was measured every 2 days at 1:00 p.m. by subcutaneously injecting mice with 15mg/mL of D-luciferin, allowing 10 min for it to access the brain, and imaging bioluminescence with 5 min exposure time. All imaging was performed using an *In Vivo* Imaging System Lumina III (IVIS, Perkin Elmer). Bioluminescence images were analyzed using Living Image Software (Perkin Elmer).

### TMZ gavage *in vivo*

TMZ (Sigma-Aldrich) was dissolved to a 50mg/mL stock solution in HPMC (Sigma-Aldrich). At the time of gavage, TMZ was diluted in 1X PBS based on mouse weight to achieve a dose of 100mg/kg with <10% HPMC. For gavage, mice were briefly anesthetized with 2% isoflurane and received between 100-200µL solution depending on mouse weight. TMZ or vehicle was administered at either ZT4 (morning) or ZT11 (evening) for 5 consecutive days after tumor growth was established at 11-13 days post-implant.

### Body weight measurements

To assess tumor burden and disease progression, mice were weighed daily starting on the day of implant. Upon 10% weight loss from pre-implant weight, mice were sacrificed in accordance with IACUC protocols.

### Mass spectrometry

Mice were anesthetized using 2% isoflurane 1 or 4hrs after receiving one dose of TMZ in the morning or evening. Blood was collected via cardiac puncture followed by decapitation and brain dissection. Tissue processing is described in Extended Materials and Methods. Ultra-high-performance LC (UHPLC)/MS/MS was performed with a Thermo Scientific Vanquish UHPLC system interfaced with a Thermo Scientific TSQ Altis Mass Spectrometer (Waltham, MA). LC/MS/MS data were processed and analyzed with the open-source Skyline software. See Extended Materials and Methods.

### Statistical Analysis

The correlation coefficient (CC) of a best-fit circadian cosine function was calculated using ChronoStar 1.0 to assess circadian rhythmicity in GBM cells (See Extended Materials and Methods). For qRT-PCR and qMSP data, JTK cycle was used to assess circadian rhythmicity. Unpaired, two-tailed Student’s t tests were used for analysis of cell proliferation and tumor size among experimental groups. Group mean differences were assessed using one-way analysis of variance (one-way ANOVA) with Tukey post hoc tests to further examine pairwise differences. A level of p < 0.05 was used to designate significant differences. All the statistical analyses were performed in Prism (version 10.0.1).

## Acknowledgements

We thank the members of the Herzog lab for discussion and comments on the manuscript. This work was supported by National Institutes of Health Grants NINDS R21NS120003 and the Washington University Siteman Cancer Center.

## Statements and Declarations

### Funding

This work was supported by the National Institutes of Health Grants NINDS R21NS120003 and the Washington University Siteman Cancer Center. Author MFGA was supported by the Washington University Neuroscience Program T32-Training Grant NIH (T32NS121881-01) and the Initiative for Maximizing Student Development (IMSD) Program Training Grant NIH (R25GM103757-10). Author ARD was supported by the National Institutes of Health National Cancer Institute (F31CA250161). Author LLT was supported by grants from Universidad Nacional de Quilmes (PUNQ 2285/22) and by Agencia Nacional de Promoción Científica y Tecnológica de Argentina (PICT 1745-2017).

### Competing Interests

The authors have no relevant financial or non-financial interests to disclose.

### Author Contributions

All authors contributed to the study conception and design. Cell experiments and analysis: M.F.G.A., L.L.T., and T.S. Animal experiments and analysis: M.F.G.A., A.R.D., L.L.T., and S.P.C.G. Mass spectrometry: K.C. The first draft of the manuscript was written by M.F.G.A., A.R.D., and E.D.H. All authors read and approved the final manuscript. M.F.G.A. and A.R.D contributed equally to this study.

### Data availability

The data generated in this study are available from the corresponding author upon reasonable request.

### Ethics approval

All vertebrate animals in this study were used in accordance with the guidelines established by the Washington University Department of Comparative Medicine (protocol 19-1136, expiration date 05/21/2023, and protocol 23-0105, expiration date 05/17/2026).

## Notes

### Competing Interest Statement

The authors have declared no competing interest.

