## Supplementarry Materials for "Circadian regulation of *MGMT* expression and promoter methylation underlies daily rhythms in TMZ sensitivity in glioblastoma"

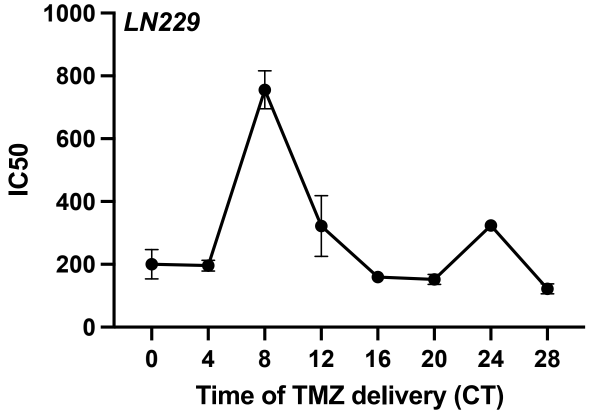


**Supplementary Fig. 1** Half-maximal inhibitory concentration (IC50) varies with circadian time of TMZ administration. TMZ administered at different circadian times *in vitro* induced LN229 cell death in a dose and time-dependent manner (n=3 per time-point each in separate cell cultures; mean ± SEM)

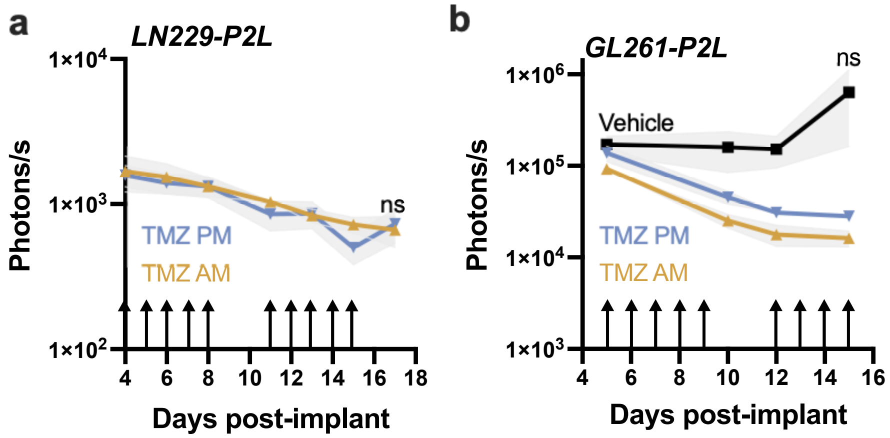


**Supplementary Fig. 2** Similar GBM sensitivity to lower TMZ dose at dawn and dusk. Separate cohorts of mice implanted with either **a)** LN229-P2L or **b)** GL261-P2L and treated daily with 70mg/kg TMZ at either 8:00am or 6:00pm (arrows) showed similar TMZ-induced tumor reduction (n=6 in AM and 6 in PM for LN229-P2L; n= 4 in AM, 4 in PM, and 4 in vehicle for GL261-P2L; mean ± SEM; ns indicates $p > 0.05$)

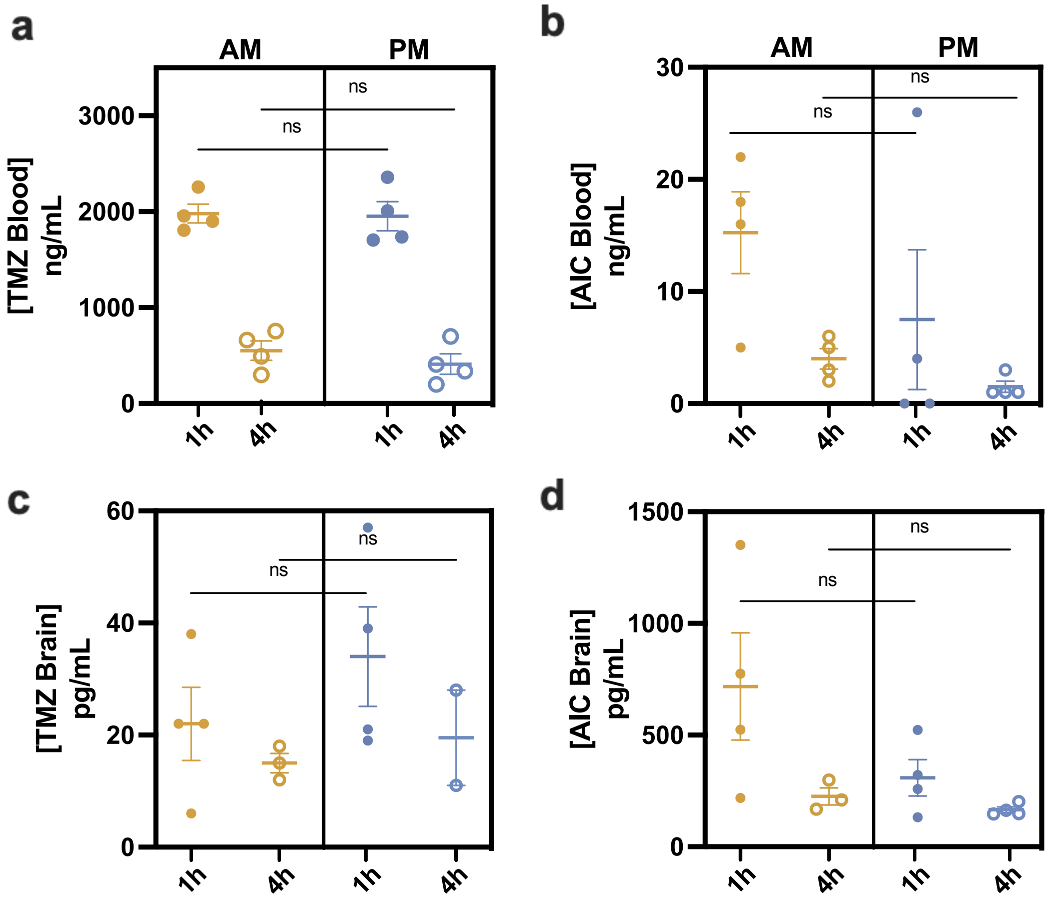


**Supplementary Fig. 3** TMZ and its metabolite AIC show similar kinetics in blood and brain when injected in the morning or evening. **a)** TMZ (*m/z* 195 → 138) and **b)** AIC (*m/z* 127 → 109) concentrations in peripheral blood showed similar decreases in levels from 1- to 4-h post-injection in the morning (ZT1, n=4) or evening (ZT11, n=4; ns indicates $p > 0.05$). TMZ and AIC concentrations in brain also did not differ with injection time (**c-d**; n=4 per group; ns indicates $p > 0.05$)

| Target | Species | Forward Primer | Reverse Primer |
|---|---|---|---|
| *MGMT* | Human | GTGATTTCTTACCAGCAATTAGCA | CTGCTGCAGACCACTCTGTG |
| *GAPDH* | Human | ATGGGGAAGGTGAAGGTCG | GGGGTCATTGATGGCAACAATA |

**Table S1**. Primers for qRT-PCR.

| Target | Species | Forward Primer | Reverse Primer |
|---|---|---|---|
| *Met-MGMT* | Human | TTTCGACGTTCGTAGGTTTTCGC | GCACTCTTCCGAAAACGAAACG |
| *Unmet-MGMT* | Human | TTTGTGTTTTGATGTTTGTAGGTTT TTGT | AACTCCACACTCTTCCAAAAAC AAAACA |

**Table S2**. Primers for q-MSP.

### Supplementary Materials and Methods

#### Glioblastoma cell culture

LN229 cells (American Type Culture Collection, CRL-261), a female human GBM cell line, were cultured in monolayer in coated T-75 flasks (Nunclon Delta coated, Fisher) using DMEM (Gibco) supplemented with 5% FBS (Fisher) and 1% Pen/Strep (Thermo Fisher). Cells were grown in a 5% CO2 incubator at 37°C. Passage number at implant ranged from six to twelve.

Glioma 261 cells (GL261, obtained from the Division of Cancer Treatment and Diagnosis Tumor Repository of the National Cancer Institute), a male murine model of GBM, were cultured in monolayer in coated T-75 flasks (Nunclon Delta coated, Fisher) using RPMI-1640 (Sigma-Aldrich) supplemented with 10% FBS (Fisher), 1% L-Glutamine (Thermo Fisher), and 1% Pen/Strep (Thermo Fisher). Cells were grown in a 5% CO2 incubator at 37°C. Passage number at implant ranged from six to twelve.

#### Cell transduction with luciferase reporters

GBM cells were transduced with lentiviral reporters expressing firefly luciferase driven by the mouse *Bmal1* (*Bmal1*-luc) (previously described in Liu et al., 2008; Zhang et al., 2009), *Period2* (*Per2*-luc) (previously described in Ramanathan et al., 2012), *Ef1α* (*Ef1α*-luc, obtained from GenTarget Inc.), or $O^6$-Methylguanine-DNA methyltransferase (*MGMT*-luc, obtained from Dr. Markus Christmann). Cells were grown in T-25cm$^2$ flasks for 24h and incubated for 10 minutes at 37ºC in 3mL complete DMEM media with 10% FBS (Thermo Fisher), 5% Pen/Strep (Thermo Fisher), and 15ug polybrene (Millipore #TR-1003-G). Following incubation, 500uL of virus stock solution was added to each culture. Media was changed after 24 h at 37ºC. Infected cells were selected using blasticidin (1.25 μg/mL, Thermo Fisher). Luciferase expression was confirmed by recording bioluminescence *in vitro*.

#### Bioluminescence recordings *in vitro*

Cells were plated in 35 mm (BD Falcon, Fisher) petri dishes supplemented with 1 mL of DMEM supplemented media containing 0.1mM D-luciferin (Goldbio), sealed with vacuum grease, and placed in a light-tight 36ºC incubator containing photo-multiplier tubes (PMTs) (Hamamatsu Photonics). Each dish was placed under one PMT and bioluminescence was recorded as photons per 180 seconds. Bioluminescence data was detrended with a 24-hour moving average and analyzed in ChronoStar 1.0.

#### *In vitro* cell growth assays and pharmacology

GBM cells were plated at a density of 80,000 cells/well in 12-well plates using one plate per time point. To assess daily rhythms in TMZ sensitivity, cells were synchronized via serum shock with 50% FBS for two hours, followed by a media change. This was done for one plate every four hours so that at one treatment time, plates spanned 0-, 4-, 8-, 12-, 16-, 20-, 24-, and 28-hours post-serum shock (HPS). At the time of treatment, cells were treated with one of four TMZ concentrations (10, 100, 200, 1000 μM) or vehicle (DMSO, 0.2%). To quantify cell numbers, we fixed cultures 72 h after treatment with cold methanol and stained cells with 4′,6-diamidino-2-phenylindole (DAPI, 2μg/mL). We imaged fluorescence with the Infinite 200 PRO plate reader (V_3.37_07/12_Infinite, Tecan Lifesciences) and transformed brightness to cell number using a linear regression equation (Y = 0.009201*X + 2153) obtained from a standard curve for each cell line. All values were normalized to vehicle to obtain a percent of cell survival.

To assess whether daily rhythms in TMZ sensitivity depend upon circadian rhythms in *MGMT* expression, cells were treated with 20μM of the MGMT inhibitor $O^6$-benzylguanine ($O^6$-BG) or vehicle (DMSO, 0.2%). For all experiments, cells were fixed after 72h with cold methanol for 10 minutes and stained with 2μg/mL of DAPI. DAPI fluorescence was measured immediately after incubation using a fluorescent plate reader (Thermo Fisher, Luminoskan), and reported as relative fluorescent units (RFU). To convert RFU to cell number, a calibration curve was performed in 12-well plates using the equation Y= 0.0096X+ 1968. Number of cells for each treatment was relativized to vehicle at each timepoint, and expressed as percent of survival.

#### Quantitative PCR (qPCR) and Methylation Specific PCR (qMSP)

RNA was extracted from 500,000 LN229 GBM cells collected every 4 hours for a period of 48 hours using TRIzol reagent. RNA was purified using the Direct-zol RNA MiniPrep Plus kit (Zymo) and cDNA was generated by RT-

PCR using SuperScript® III First-Strand Synthesis System (Thermo Fisher). For qMSP, bisulfite DNA conversion was performed with the EZ DNA Methylation-Gold kit following their standard protocol. Gene expression changes were further probed using iTaq™ Universal SYBR® Green Supermix (Bio-Rad). The primer sequences for qRT-PCR are listed in Table S1 and for qMSP in Table S2, and were obtained from previous publications (Yoshioka M., et al. (2018), Kreth S., et al. (2011)). PCR amplification was carried out at 40 cycles with 10 ng of template DNA in triplicates. Protocol is as follows: Cycle 1: 95ºC for 3 min; Cycle 2: 95ºC for 30 sec; Cycle 3: 60ºC for 30 sec; repeat step 2-3 for 39 more times; Cycle 4: 72ºC for 1 min. Negative controls included no reverse transcriptase reactions and no template DNA samples. All procedures were done in triplicate in two biological replicates.

**TMZ gavage *in vivo***
TMZ (Sigma-Aldrich) was dissolved to a 50mg/mL stock solution in HPMC (Sigma-Aldrich). At the time of gavage, TMZ was diluted in 1X PBS based on mouse weight to achieve a dose of 100mg/kg with <10% HPMC. A 25mm, 22G, stainless steel, round bulb gavage bent needle was used for gavage. When not in use, needles were stored in 70% ethanol. For gavage, mice were briefly anesthetized with 2% isoflurane and received between 100-200μL solution depending on mouse weight. TMZ or vehicle was administered at either ZT4 (morning) or ZT11 (evening) for 5 consecutive days after tumor growth was established at 11-13 days post-implant.

**Mass spectrometry**
*Blood collection*
Mice were anesthetized using 2% isoflurane, collected 200uL blood via cardiac puncture followed by decapitation and brain dissection. Blood was immediately mixed with 10% 0.5M sterile EDTA (Sigma) and centrifuged (15 min at 2,000 x g). Plasma was stored at -80°C until use. Upon thawing, 100ng/mL dacarbazine (Selleck Chemicals) was added to the plasma sample in a 2mL Eppendorf tube and split it into two 50uL aliquots. We acidified one aliquot with 10µL of 1 N HCl (Fisher) and alkalified the other with 10uL of 1N NaOH (Fisher). Proteins were precipitated using 100µL of cold acetonitrile (Thermo Scientific), then vortexed and centrifuged (10 min at 5,000 rpm) samples. 95uL were transferred to LC vials for analysis. Samples were stored at -80°C until analysis.

*Brain collection*
Following decapitation, whole brains were dissected and washed briefly in 1x PBS. We snap froze brains in liquid nitrogen and stored them at -80°C until extraction. For extraction, we weighed snap frozen tissue and obtained a section of cortex (28-40mg). We ground samples with a plastic homogenizer tube (Fisher). For each 1mg sample we added 40uL 9:1 methanol: water and 100ng/ml dacarbazine. We vortexed (30 sec), submerged in liquid nitrogen (1mn), thawed (10 sec), and bath sonicated at 25°C (10mn), all samples four times. We acidified 100uL sample with 20uL 1N HCl. We alkalified 100uL sample with 20μL 1N NaOH. Samples were stored overnight at -20ºC to complete precipitation of proteins. We centrifuged samples (10 min at 14,000 rpm at 4ºC) and transferred 100μL supernatant to LC vials for analysis.

*LS/MS/MS*
Ultra-high-performance LC (UHPLC)/MS/MS was performed with a Thermo Scientific Vanquish UHPLC system interfaced with a Thermo Scientific TSQ Altis Mass Spectrometer (Waltham, MA). Reversed-phase liquid chromatography (RPLC) separation was accomplished by using an Acquity UPLC HSS T3 column (150 mm x 2.1 mm, 1.8 μm, Waters). The column compartment temperature was 40°C, and the flow rate was set to 300 $\mu L*min^{-1}$. Mobile-phase solvents were composed of A = 0.5 mM ammonium formate, 0.1% formic acid in water and B =100% methanol. The following linear gradient was applied: 0 – 1 min, 5% B; 1 – 5 min, 90% B; 5 – 6 min, 90% B, 7 min, 5% followed by a re-equilibration phase of 4 min at 300 $\mu L*min^{-1}$. The samples were kept at 4°C in the autosampler. The injection volume was 5 μL for all samples.

Data were collected with the follow settings: sheath gas flow 50 Arb, auxiliary gas flow 10 Arg, sweep gas flow 1 Arb, ion transfer tube temperature 325°C, and vaporizer temperature 350°C. The spray voltage was 3.5 kV for positive polarity. A cycle time of 0.8 s was used. The Q1 and Q3 resolutions were 0.7 and 1.2 Da, respectively. Multiple reaction monitoring (MRM) was performed to detect *m/z* 195 → 137 for TMZ and *m/z* 127 → 109 for AIC. For the CID gas pressure, 1.5 mTorr was used. LC/MS/MS data were processed and analyzed with the open-source Skyline software.

**Statistical Analysis**

Circadian $R^2$ value (CC) was calculated using ChronoStar 1.0 for *in vitro* bioluminescence traces. This statistical method measures how well a cosine curve fits a time series and is used to assess circadian rhythmicity. CC values above 0.7 were considered to be circadian. Unpaired, two-tailed Student's t tests were used for analysis of cell proliferation and tumor size among experimental groups, and changes in GBM proliferation index *in vivo*. Group mean differences were assessed using one-way analysis of variance (one-way ANOVA) with Tukey post hoc tests to further examine pairwise differences. A level of $p < 0.05$ was used to designate significant differences. All the statistical analyses were performed in Prism (version 10.0.1).
